# mRNA codon optimization on quantum computers

**DOI:** 10.1101/2021.02.19.431999

**Authors:** Dillion M. Fox, Kim M. Branson, Ross C. Walker

## Abstract

Reverse translation of polypeptide sequences to expressible mRNA constructs is a NP-hard combinatorial optimization problem. Each amino acid in the protein sequence can be represented by as many as six codons, and the process of selecting the combination that maximizes probability of expression is termed codon optimization. This work investigates the potential impact of leveraging quantum computing technology for codon optimization. An adiabatic quantum computer (AQC) is compared to a standard genetic algorithm (GA) programmed with the same objective function. The AQC is found to be competitive in identifying optimal solutions and future generations of AQCs may be able to outperform classical GAs. The utility of gate-based systems is also evaluated using a simulator resulting in the finding that while current generations of devices lack the hardware requirements, in terms of both qubit count and connectivity, to solve realistic problems, future generation devices may be highly efficient.

## Introduction

Protein sequences can be encoded by an enormous multitude of possible nucleotide sequences. The degenerate mapping between amino acids and synonymous codons entails an exponential relationship between the number of potential nucleotide sequences and the length of the polypeptide chain. However, different nucleotide sequences encoding the same protein may exhibit dramatically different outcomes in expression systems.^1–4^ Furthermore, recent studies have shown that codon selection can impact downstream processes such as protein folding and function,^1–3^ which is particularly important for use-cases such as recombinant protein therapies.^5^

Codon optimization is a procedure designed to increase gene expression based on a heuristic scoring function^6^ with many scoring functions having been proposed.^7–10^ Some of the more common scoring functions seek to optimize the fraction of G and C bases,^11–15^ match the codon usage bias of the host expression system,^16–21^ and/or attempt to disrupt the formation of mRNA secondary structure.^7,17,22^ The vast solution space is most commonly sampled using genetic algorithms (GA) that seek to evolve solutions by introducing synonymous codon mutations and propagating favorable substitutions through generations.^12,16,20,23–25^ However, other methods have been proposed.^18,26,27^ While classical approaches such as GAs can be highly performant, the fraction of solution space that is sampled in a fixed number of iterations decreases exponentially as the polypeptide chain length grows. Thorough sampling of the solutions space is therefore often intractable with biologically relevant use-cases. In this study we investigate the viability of programming quantum computers to efficiently identify high-quality solutions scored with arbitrary objective functions.

Recent advances in quantum information science and technology have elucidated the potential for quantum devices to outperform classical devices in a narrow range of applications.^28,29^ Among these, certain types of combinatorial optimization problems are among the most promising for near-term advantage.^30^ The field of quantum computing is developing rapidly in terms of both hardware and algorithms. Each type of quantum computing technology offers a unique set of strengths and weaknesses, and applications are often tailored to compliment the strengths of each device. While there are many physical realizations of quantum technologies, we focus on two markedly different models; adiabatic quantum computers (AQC) and gate-based quantum computers.

AQCs, sometimes referred to as quantum annealers, are most commonly used to solve high-dimensional combinatorial optimization problems.^31–33^ For example, a recent study showed that protein design, which requires combinatorial optimization of rotomeric states, can be accelerated with an AQC.^34^ Current implementations of AQCs offer nearly two orders of magnitude more qubits than state of the art gate-based devices, offering the potential to address realistic sized problems. The general class of AQC algorithms are classified as metaheuristic methods for solving local optimization problems in multivariate spaces.^31^ These approaches are similar to simulated annealing but exploit the phenomena of quantum tunneling, instead of thermal activation, to hop out of local energetic minima.^35^

An alternative quantum computing technology, and the technology that was recently used to demonstrate quantum supremacy,^28,29^ centers on gate-based instructions. Current generations of non-error corrected hardware, termed Noisy Intermediate Scale Quantum (NISQ),^36^ can be programmed to solve combinatorial optimization problems using variational methods such as the Quantum Approximate Optimization Algorithm (QAOA).^37^ Gate based quantum computers are presently, however, less mature than AQCs and even the most capable devices to date lack the number of qubits and connections between qubits needed to solve realistic combinatorial optimization problems. As the technology matures however there is an expectation that qubit count and connections between qubits will improve substantially. There is also the expectation that error rates will decrease, and general-purpose error correction may become possible. Developing and testing suitable algorithms ahead of the technology development curve is thus a worthwhile endeavor and it is therefore common practice to utilize simulators, running on classical computing hardware, to permit evaluation of quantum algorithmic functions in the absence of real-world hardware to evaluate performance on.

Designing novel algorithms to execute on quantum devices requires deep expert knowledge of the devices, quantum information science, and quantum software stacks. However, there are a few classes of ubiquitous problems that are readily solvable using tools built into many widely available software packages. The Binary Quadratic Model (BQM) is perhaps the most archetypal example and can be found at the core of many familiar problems such as the Ising model.^38,39^ The energy of a BQM can be described by a Hamiltonian with the general form

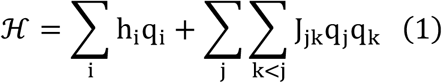

where q_i_, q_j_, and q_k_ represent the values of the qubits, which can either be {0, 1} or {−1, 1} for binary or spin representations, respectively, h_i_ are the one-body terms, and J_jk_ are the two-body interactions. For the Ising model, h represents the physical spins and J represents the energy of the interactions between the spins.

The methods section below shows that the codon optimization problem can be mathematically formulated as a BQM and thus implemented on a variety of competing quantum computing platforms. This representation requires translating the scoring function used in traditional approaches into a quadratic Hamiltonian where the eigenstates represent nucleotide sequences and the eigenenergies represent the scores. Implementing this program on the D-Wave Advantage 1.1 AQC shows that it identifies high quality solutions that are competitive with a basic implementation of a genetic algorithm programmed with an equivalent scoring function. Implementing a version of this program for IBM Q devices, while successful, shows that modelling practical systems requires too many qubits to be run on even the most advanced gate-based devices available (e.g. IBM’s 65-qubit Hummingbird device).^40^ However, executing the model on an IBM noisy simulator^41^ demonstrates the potential of the algorithm. Finally, we comment on the potential usability of the BQM approach on current and future generations of quantum computers.

## Results

Codon optimization was implemented as a BQM on quantum devices with a Hamiltonian designed to optimize GC-content, minimize sequentially repeated nucleotides, and optimize codon-usage bias (see equation (15) in Methods for details). Quantum devices use qubits to store data, which decode digitally to 0’s and 1’s upon measurement, but which also may be in a superposition of 0 and 1 during the calculation. To encode classical genetic data into a quantum device, every possible codon that can map to the target polypeptide sequence is required to be explicitly represented by a physical qubit. The qubits which return “1” upon measurement represent the codons selected at each position in the polypeptide sequence. Therefore, only 1 qubit (codon) for each position in the polypeptide sequence can be in a “1” state, and the rest must return “0” upon measurement (Figure 1a). This scheme is enforced by constructing a 2-body penalty matrix which adds infinite energy to pairs of codons that map to the same position in the polypeptide sequence (Figure 1b). The final sequence is determined by recording the values of the qubits and concatenating the corresponding codons of the qubits in the “1” state (Figure 1c).

**Figure 1.**
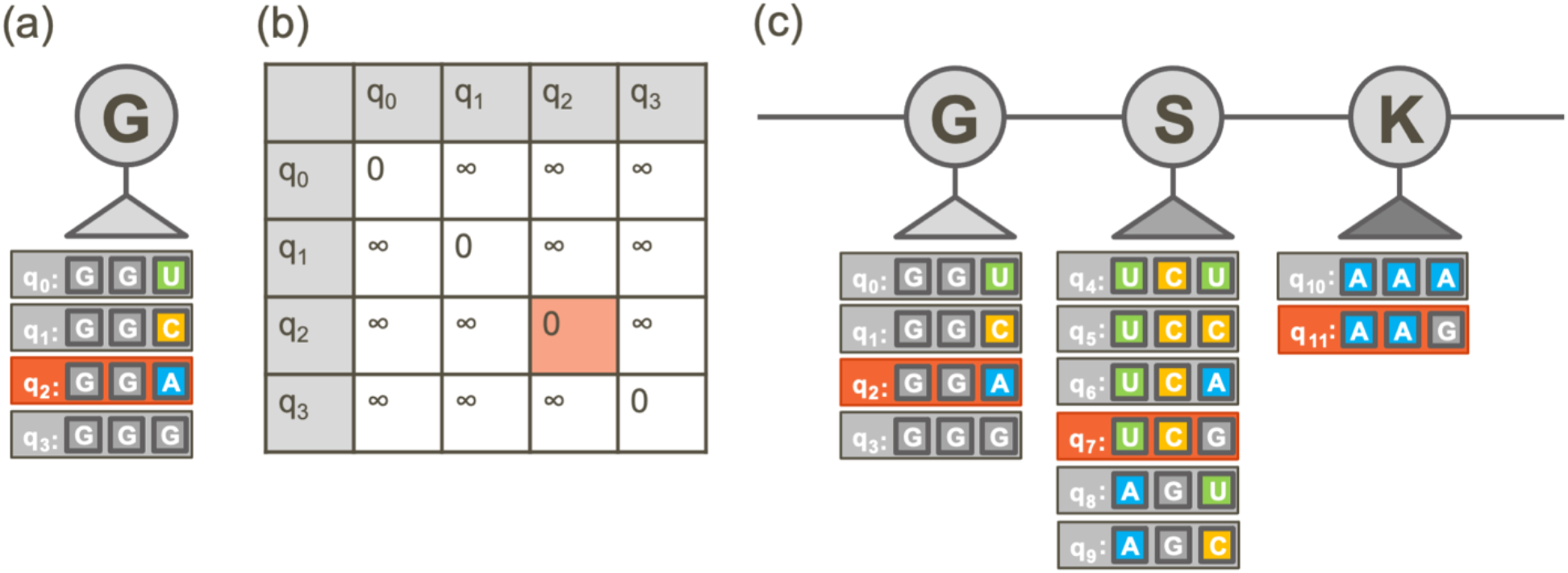
(a) Example mapping of each possible codon for the amino acid Glycine (gray oval labeled “G”) to qubits. Gray boxes represent qubits labeled q_0_…q_3_, and the codon assigned to each qubit is shown next to the qubit label. (b) A penalty matrix is constructed to add infinite energy to cases where more than one codon is in the “1” state. In this example, qubit q_2_ is in the “1” state and the rest are in the “0” state, which returns an energetic penalty equal to 0. (c) Example mapping of codons for protein sequence GSK… to qubits. One codon is selected for each position in the sequence, highlighted in orange.

### AQC accuracy and quality of scores

The goal of the optimization is to find the combination of codons that minimizes the Hamiltonian (or the objective function for the GA). In theory, AQCs should be able to find the ground state of the input Hamiltonian. However, due to thermal fluctuations and limited quantum processing unit (QPU) time, low energy solutions that are near, but not equal to the ground state are expected. Furthermore, as the size of the problem increases, the probability of annealing to an optimal eigenvalue decreases. See the “BQM challenges and limitations” section in the Supplementary Information for further discussion of technical challenges specific to heavily constrained problems such as codon optimization.

While it is not possible to calculate the true ground state for large problems, comparisons can be made to other approximate methods, e.g., GA approaches, to contextualize the results and performance of the AQC approach. Additionally, D-Wave offers hybrid solvers that augment the AQC with classical methods to optimize the results. Direct programming of the AQC yielded excessively noisy results with high variance in the estimation of the ground state. The hybrid solver was thus selected to more reliably optimize the input Hamiltonian.

#### Peptides of length 20

The baseline performance of the AQC implementation of codon optimization was evaluated using 63 peptide fragments of length 20 derived from the human severe acute respiratory syndrome coronavirus 2 (2019-nCoV, SARS-CoV-2) spike glycoprotein sequence (UniProtKB–P0DTC2) and compared with a conventional single-threaded GA implementation. See the Genetic Algorithm Validation section in the Supplementary Information for performance metrics. The results of running the AQC and the GA 20 times each are reported in Figure 2. There is an approximately linear relationship between the optimization scores, with an average ratio close to 1:1. The minimum eigenvalue identified by the AQC matched the minimum score obtained by the GA to machine precision in 77% of the peptide fragments that were considered. In the remaining cases, the GA identified a lower score than the AQC. The AQC was programmed to run for a total of 6.0 s including preprocessing and communication between the classical and quantum components. The average execution time for the GA was 1.09 s with a standard deviation of 0.06 s.

**Figure 2.**
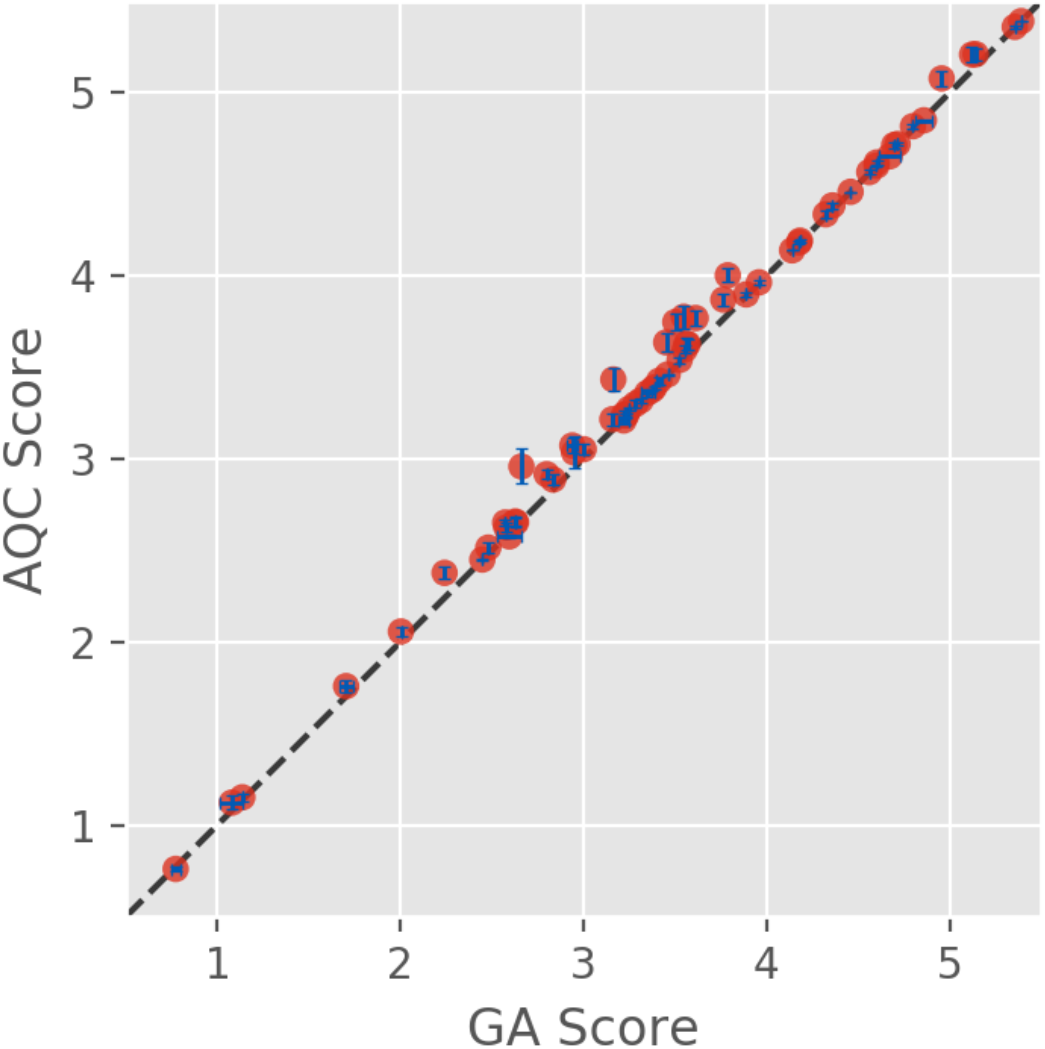
Eigenvalues measured by AQC vs GA scores. Lower scores indicate higher probability of expression. Dashed line represents y=x. Error bars represent the standard deviation of 20 trials and are shown in blue.

#### Full-length proteins

The Leap Hybrid solver is capable of solving codon optimization problems expressed as a BQM with up to ∼1,000 amino acids. A selection of full-length sequences (see Test Applications in Methods) was run on both the AQC and the GA (Figure 3). Each sequence was allotted 50 s of compute time on the AQC. The GA was run for 6000 generations, determined through testing to be the point at which the systems asymptotically converged to a solution (see Genetic Algorithm Validation in the Supplementary Information). The average execution time for the GA was 10.6 minutes with a standard deviation of 0.9 minutes.

**Figure 3.**
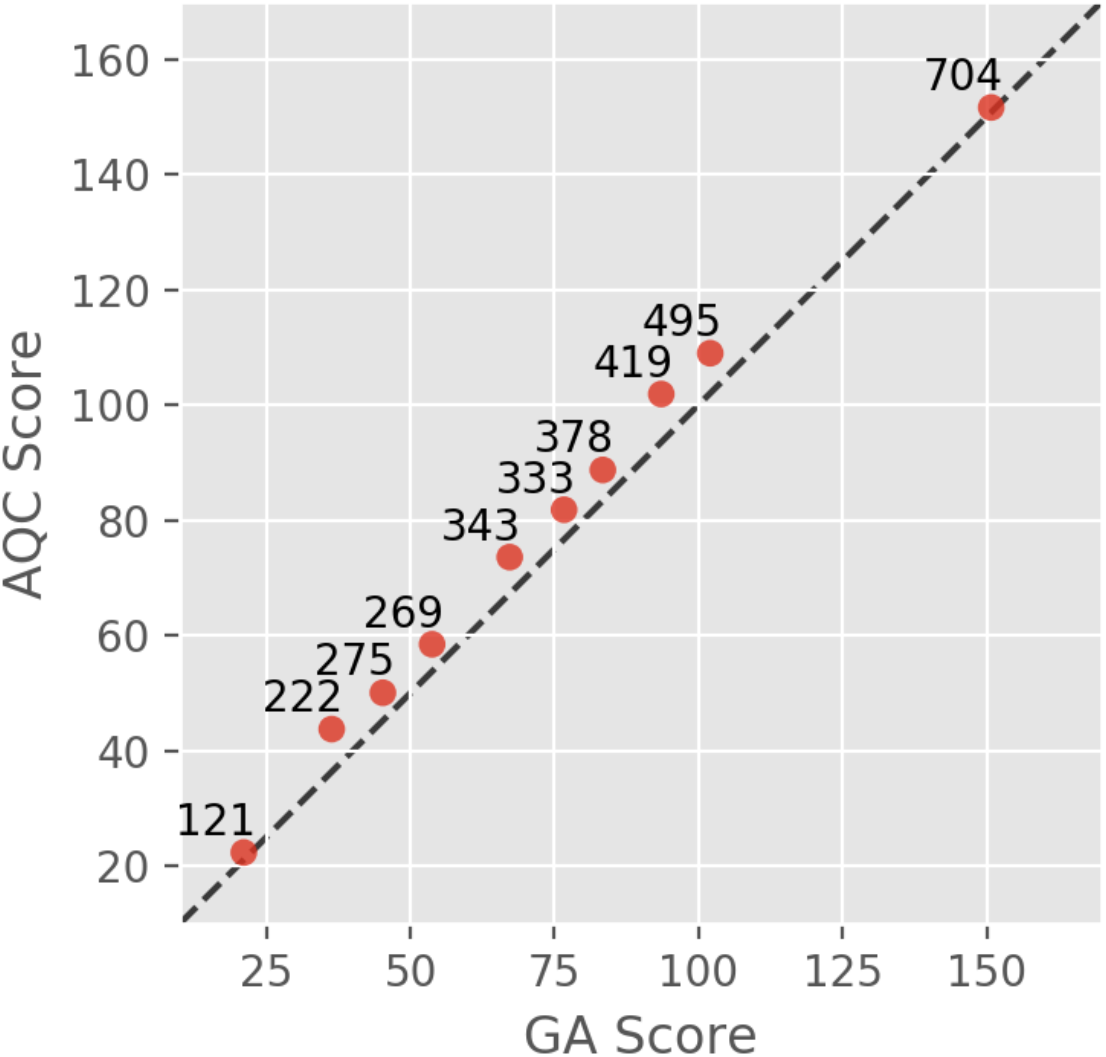
AQC vs GA for 10 full-length proteins. Dashed line represents y=x. Lower scores indicate higher probability of expression. Point labels indicate number of amino acids.

#### Relationship between scoring function and scalability on quantum hardware

The choice of scoring function can have a dramatic impact on the size of the model. For example, the Hamiltonian representing GC-content requires explicit calculation of all off-diagonal elements of the two-body interaction matrix. In other words, all logical qubits must be fully connected. The D-Wave Advantage system is state-of-the-art in terms of number of qubits and connectivity, described by a P_16_ Pegasus graph,^42^ and the best minor embedding scheme for a fully connected graph of logical qubits was heuristically found to support a maximum of 180 nodes when interfacing directly with the QPU. However, there are two-body Hamiltonians that do not require couplers between all logical qubits. The Hamiltonian penalizing sequentially repeated nucleotides only requires couplers between qubits mapping to neighboring sequence positions. In this case, the task of minor embedding is straightforward and would only require a few additional physical qubits to represent the required logical qubits on the D-Wave Advantage system.

### Gate-based simulator results

The gate-based approach was simulated using the IBM Qasm noisy simulator, which can simulate up to 24 fully connected qubits. The BQM implementation was identical to the scheme described by the AQC (Figure 1), and the optimization was carried out using QAOA.^37^ The codon optimization simulation was run on 313 peptides of length four and one peptide of length three, each requiring between 7-24 logical qubits with full connectivity. In the most resource-intensive scenario, there could be four amino acids in a row that each map to six qubits, and the algorithm is tasked with finding the ground state out of 6^4^ = 1296 possible states using 24 qubits (the maximum supported by the simulator). The task is therefore small enough to compute the exact result with the NumPyMinimumEigensolver exact solver as a comparison to the simulation.

The simulated scores vs the exact scores are shown in Figure 4a. The quantum algorithm identified the exact solution for 59 peptides, and overall, 84% of the trials yielded valid results. However, there were 51 cases where the quantum algorithm returned results which had an invalid mapping between codons and qubits. In each of the invalid trials there was at least one amino acid position lacking a selected codon.

**Figure 4.**
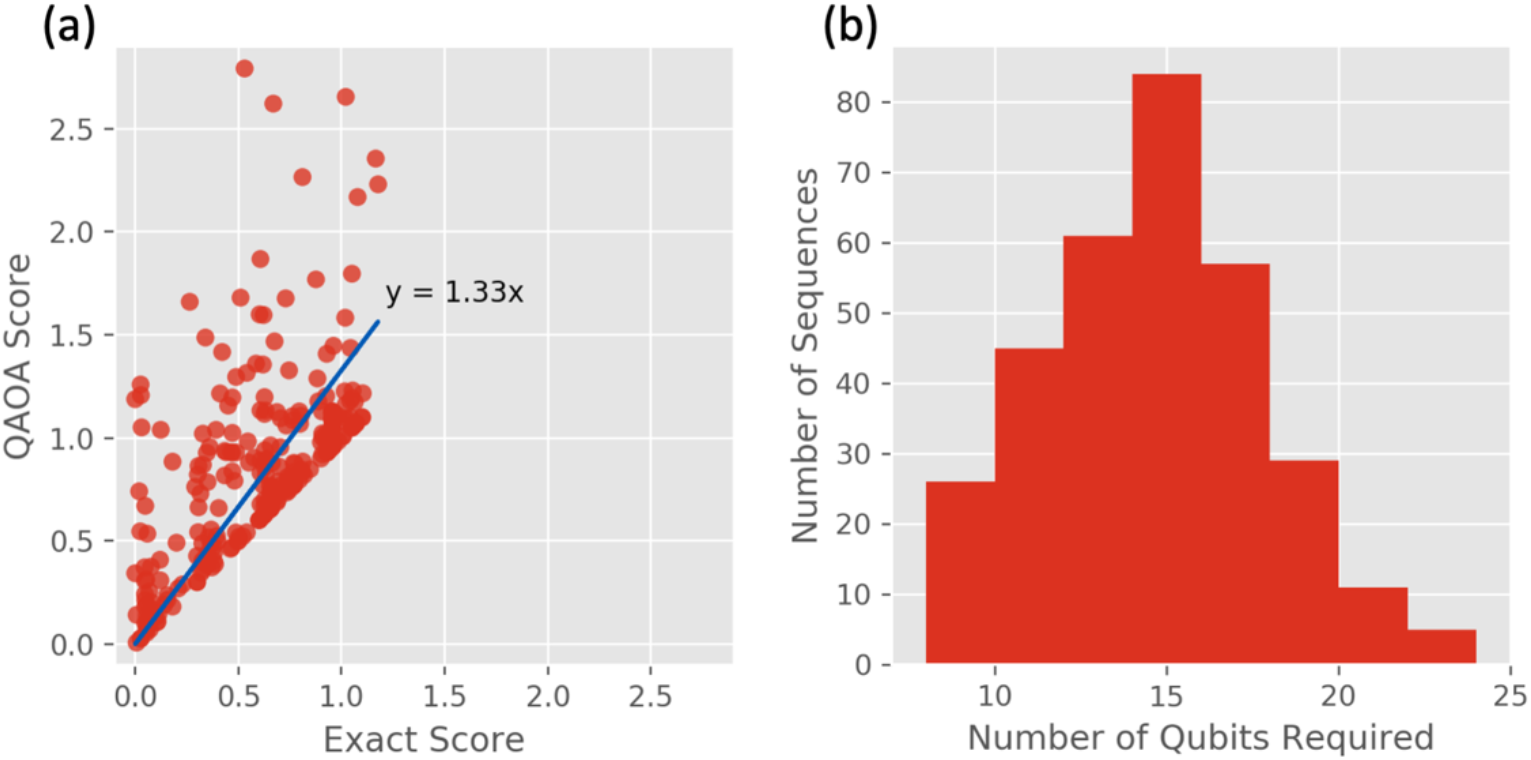
(a) QAOA simulated scores vs exact scores. Each point represents a single run of the simulator for each peptide fragment. Lower scores indicate higher quality results. Trials that returned solutions with invalid mapping between codons and amino acids were excluded. Line of best fit, displayed in blue, shows on average scores from the quantum algorithm are 33% higher than the classical algorithm. (b) Number of qubits required to simulate each of the 313 peptide fragments of length 4.

This type of error was more common in cases where more qubits were required to run the simulation (minimum 16 qubits), and therefore the optimization task was more difficult. The success of the calculation was dependent on the random seed used for the noisy simulator; rerunning trials that failed to return valid results with different random seeds changed the outcome and, in each case, a valid result was eventually found. The instability of the simulation and its dependence on the number of qubits required to run the simulation is an artifact not just of the simulated noise but also an inherent consequence of the variational algorithm employed by the simulator, which is not guaranteed to converge every time. See the “BQM challenges and limitations” section in the Supplementary Information for discussion about the exponentially small subset of valid solutions in the total possible solution space as the number of required qubits is increased. These results imply that it may be impossible to effectively optimize polypeptides of biological relevance on NISQ devices. Further studies are required to determine if this behavior could be improved by making use of alternative variational optimization algorithms or whether fundamental improvements in error correction are needed at the hardware level.

## Discussion

Codon optimization is a classic example of a biological problem with exponential scaling in solution space. While there exist classical machine learning and artificial intelligence methods that improve sampling, quantum technology may offer an alternative or complimentary approach to enhance the ability to identify optimal samples from these distributions. Current generations of quantum hardware are mature enough to test ideas for novel algorithmic approaches to problems in life sciences but are not yet capable of outperforming classical devices. In this study codon optimization is reformulated to be readily implementable on quantum devices and the viability of the method is demonstrated on both adiabatic and gate-based quantum computers.

The D-Wave Systems Advantage 1.1 system was heuristically determined to be capable of solving codon optimization problems programmed directly into the QPU up to approximately 180 codons with scoring functions requiring all logical qubits to be fully connected, which maps to 30 amino acids in the most resource intensive cases. If a scoring function is selected that does not require full connectivity, such as the Hamiltonian describing repeated nucleotides, then nearly all of the logical qubits could map directly to physical qubits and the system size could theoretically be scaled to accommodate sequences of up to ∼1,000 amino acids. There is likely significant room for performance improvements in the quantum, hybrid, and classical methods. As AQC hardware matures it will be possible to address questions of scalability for larger sequences and critically assess the potential for quantum technology to surpass classical techniques.

The IBM Experience provides free access to small quantum devices and noisy simulators. The largest quantum device freely available, Melbourne, contains 15 qubits with 2-3 couplers between them. The BQM presented in this study requires full connectivity between the logical qubits. Melbourne is therefore able to represent a select set of 2-amino acid systems in which each amino acid maps to one or two codons. Meanwhile the IBM Qasm simulator allows up to 24 fully connected simulated qubits, which is generally limited to 4 amino acids. This simulation approach is sufficient to use QAOA to solve small proof of concept problems, but realistically thousands of qubits with high connectivity are required to run biologically relevant sequences. Given IBM’s current public Roadmap for Scaling Quantum Technology,^40^ devices with this capacity are not expected to be available to the public before 2024. Utilizing the simulator, the QAOA algorithm had variable performance, identifying the true ground state solution in nearly 20% of the trials, and failing to identify a valid solution in 16% of the trials. As noted above the success of the calculation was dependent on the random seed used for the noisy simulator; valid results for each failed run can be obtained by rerunning with different random seeds until a valid result is found. Further investigation on physical devices is required to determine the limit of the accuracy and precision of the method.

There is a vast body of literature discussing codon optimization techniques for protein expression. This study contributes to the field by offering a novel approach to sampling the vast solution space with an emerging technology. The considerations that went into the construction of the Hamiltonian were designed to highlight some of the implementational nuances associated with common scoring functions. For example, the expression for measuring optimality of GC-content requires full connectivity between qubits. However, counting repeated nucleotides between neighbors only requires some qubits to be coupled. Codon usage bias can be factored into the Hamiltonian using one-body terms, but one could imagine a more thorough approach which compares the distribution of codons selected in the sequence compared to the reference distribution, which would require two-body terms. Similarly, implementing the popular “one amino acid-one codon” strategy would require codons from each type of amino acid to be coupled,^18,21,43^ adding many two-body terms but not as many as optimization of GC-content. Thus, the particular Hamiltonian studied here serves as a demonstration of the efficacy of the method at evaluating a realistically complicated objective function.

Further studies are required to evaluate these samples beyond simple numerical scores to determine the value this approach could add to the field of protein expression using future generations of quantum hardware. Current quantum hardware is subject to high levels of noise and is therefore not competitive with classical techniques in most practical applications. However, quantum hardware and techniques are advancing exponentially in terms of both scale and tolerance to noise. This rapid advancement coupled with the expectation that scaling with problem size is significantly reduced on quantum hardware compared with classical methods implies that future quantum approaches could provide a significant performance advantage for optimization of NP problems in life sciences.

## Methods

### Codon optimization algorithm

Classical scoring functions can be reinterpreted as a Hamiltonian by separating one- and two-body interaction terms. There is considerable research and ongoing debate of the proper way to score nucleotide sequences for expression,^5,8,18–21,24–27,43,10–17^ as well as arguments against the use of codon optimization in some cases.^5^ The purpose of this study is not to contribute to the discussion of whether or not codon optimization is an appropriate tool for any given context or the optimal way in which it should be performed but rather to investigate a novel method of sampling the vast solution space. The optimization task is therefore restricted to three considerations that capture simple countable properties from the sequences themselves. This includes discussion of how to formulate energetic terms in the Hamiltonian that serve to minimize:

1. Codon usage bias.
2. The difference between GC-content and a target value.
3. The number of sequentially repeated nucleotides.

Energetic terms can be combined into one Hamiltonian or used on their own, just like a scoring function in a GA. Furthermore, the Hamiltonian could be extended using any type of scoring function that can be broken down into one- and two-body interaction terms. The following sections outline the mathematical representations of these optimization tasks and show how they naturally map to a Hamiltonian with a form compatible with a BQM.

Two additional constraints are imposed to add energetic penalties to combinations of codons that do not translate to the query sequence. The first constraint adds a small linear shift to the one-body term of each qubit. Shifting the potentials increases the energetic favorability of including more codons in the sequence. Similarly, the other constraint adds a significant energetic penalty to codons mapping to the same position in the amino acid sequence. The combination of these two potentials optimizes the energetic score of valid combinations of codons compared to invalid combinations.

#### Incorporating codon usage bias

Codon usage frequency varies by host system.^44^ Therefore, the scoring function is tailored to match the expression system. For this study, codon usage frequencies for e. coli are imported from the python-codon-tables 0.1.10 library.^45^ Let *C_i_* represent the frequency of finding codon *C* at position *i*. The potential is thus designed to return a large penalty for rare codons (where *C_i_* is small) and incur a negligible penalty for codons readily available to the system (where *C_i_* is large). One such function is the log of the inverse multiplied by −1. This function yields the desired behavior of adding large penalties to rare codons and adding small penalties to accessible codons. However, the function is undefined at *C_i_* = 0. If a host system truly does not have access to a given codon, then any sequence containing the codon is not expressible, and the probability of expression would be zero. However, the scoring function must be restricted to finite decimal values, so an infinitesimal value, *ε*_*f*_, is added to the denominator to avoid undefined values. For a system containing *N* possible codons, the Hamiltonian is given by:

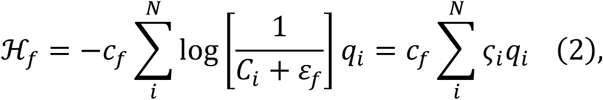

where *c*_*f*_ is a tunable constant, *q*_*i*_ ∈ {0,1} is the value of the qubit, and ***ζ*** is a vector containing the values of the log inverse codon usage frequencies. Given the binary *q*_*i*_ values, the Hamiltonian only penalizes codons that are “selected”, represented by qubits with value *q*_*i*_ = 1.

#### Optimize target GC concentration

To optimize the GC concentration of a nucleotide sequence, ρGC, a cost function *Δ* must be introduced to minimize the difference between *ρGC* and the target GC concentration, *ρT*. The simplest objective function satisfying this constraint is a quadratic function,

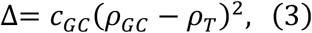

where *cGC* is a tunable constant. The GC content is calculated by summing the number of G’s and C’s in the sequence of length *N* and normalizing by the number of nucleotides in the sequence

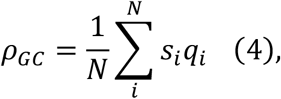

where *si* is an integer representing the number of G’s and C’s in codon *i*, and *qi* represents the value of qubit *i*. By expanding equation (3), a form similar to a Binary Quadratic Model (BQM) formulation becomes apparent:

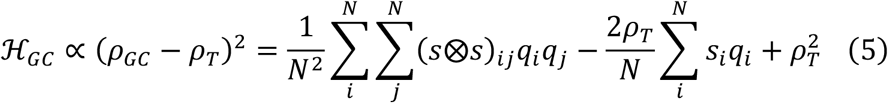

The matrix represented in the double sum needs to be restricted to a sum over the upper triangular elements, consistent with equation (1). By decomposing the sum into the trace and a term that sums the contributions of the off-diagonal elements, the sum can be restricted to the upper triangular elements. The trace requires a single summation over *Si^2^*. Since qubits map to binary values, they are idempotent with themselves and therefore *qi^2^*=*qi*.

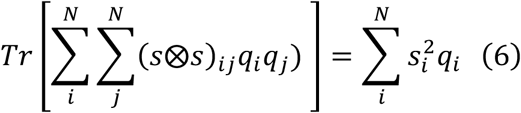

Since the matrix is symmetric, all off-diagonal terms are accounted for in a upper triangular form by multiplying by 2. Thus,

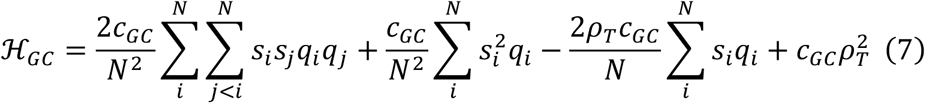

recapitulates the quadratic cost function (equation (3)) in a form consistent with a BQM (equation (1)). See the GC-Content Derivation section in the Supplementary Information for a full derivation.

#### Minimize sequentially repeated nucleotides

To minimize the number of repeated nucleotides in a sequence, all codons mapping to sequential positions in the amino acid sequence are compared. Let *r(C*_*i*_, *C*_*j*_) represent a quadratic function that returns the maximum number of repeated sequential nucleotides between codons *C_i_* and *C*_*j*_, shifted to the origin for null cases by subtracting one. For example,

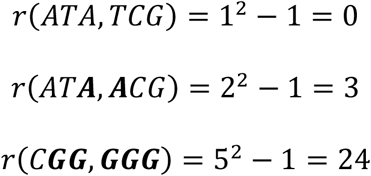

Repeated nucleotide penalties are stored in a matrix ***R***,

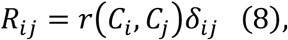

where the delta-function returns 1 if the codons map to sequential positions and 0 otherwise.

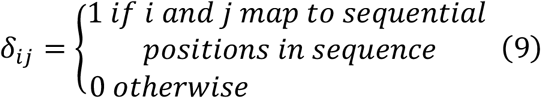

Supplementary Figure S1 shows the result of applying this function to the system in Figure 1. The total repeated nucleotide penalty for a given nucleotide sequence is given by

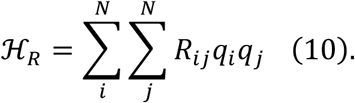

Matrix ***R*** is upper triangular, so the pairwise sum can be restricted to upper triangular elements without changing the result, making it compatible with the BQM model (equation (1)). A tunable constant, *c_r_*, is introduced to weight the contribution in the Hamiltonian, resulting in the form:

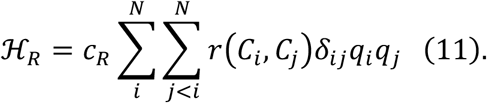

#### Additional constraints to the Hamiltonian

The energetic terms described above are designed to add penalties to particular sequence properties. The ground state energy of this type of objective function is zero since introducing codons increases the score. To counteract this tendency, a constant factor, ε, is subtracted from the one-body term of each codon, thereby increasing the energetic favorability of introducing codons to the system. The absolute value of the constant must exceed the largest value in the one-body vector *h* (equation (1)).

To negate the possibility of assigning more than one codon to a given position, a site-specific delta-function is introduced that applies an effectively infinite penalty to pairs of codons assigned to the same position.

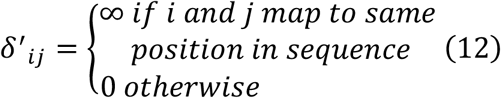

Supplementary Figure S2 shows the result of applying equation (12) to the example system referenced in Figure 1. For a system with *N* possible codons, the Hamiltonian is modified by adding the following terms:

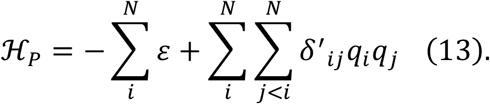

#### Implementation of objective function

The Hamiltonian representing the total “energy” of a nucleotide sequence is computed by summing the contributions of the expressions defined in the previous sections.

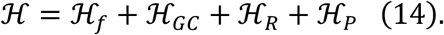

Expanding and rearranging the terms gives the form in equation (15):

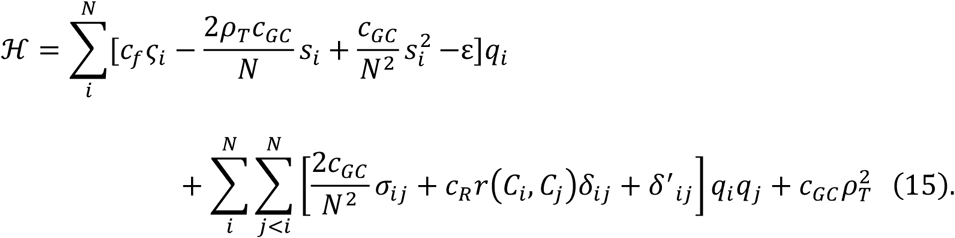

This form is consistent with the BQM formalism (equation (1)) and can be directly implemented into BQM frameworks.

### Algorithm implementations

The current approach to performing calculations on quantum devices requires the interaction terms to be precomputed on classical devices and read into the quantum devices via specialized APIs. The one- and two-body interaction terms from equation (15) were precomputed in python 3.7 using standard libraries and numpy^46^ arrays. The numpy arrays were converted to dictionaries in accordance with the expected input for the quantum device libraries.

The execution of the BQM was carried out using libraries described in the following sections. Each calculation was run 20 times for small peptide fragments (<100 residues), and full-length proteins were only run one time due to QPU resource constraints. Table 1 provides the values of the constants used in the objective function. The eigenstate returned by the quantum devices was converted back to a nucleotide sequence and translated to a polypeptide sequence to verify the validity of the result.

**Table 1.**
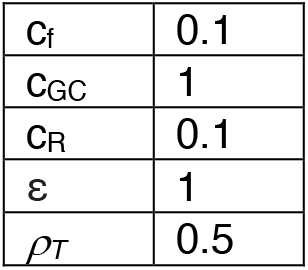
Constants used in equation (15).

Furthermore, the nucleotide sequence was scored using the classical scoring function used by the GA, and the result was compared to the eigenvalue returned by the quantum devices. Eigenvalues and classical scores agreed to 8 decimal places.

#### D-Wave Advantage 1.1

The codon optimization BQM was implemented on the D-Wave Advantage System 1.1 utilizing the Leap Hybrid Solver^1^. This adiabatic quantum device contains more than 5,000 superconducting qubits. Each qubit is connected to 15 others described by a Pegasus P_16_ graph.^42^ The Advantage system was accessed through the D-Wave Leap web interface, which serves as an access point to QPU hardware as well as an integrated developer environment with built-in support for the full D-Wave API.

The program was constructed and executed using python libraries provided by D-Wave systems. The BinaryQuadraticModel class in the dimod 0.9.10 python library was used to construct the model from the classically prepared data and convert it to a data structure compatible with the quantum device. The one- and two-body interaction terms were precomputed, stored in numpy arrays, and passed into the BinaryQuadraticModel instance along with an offset of 0.0 and the dimod.BINARY representation. The model was executed using the LeapHybridSampler classes in the dwave.system python library. The solver was allotted 6 s of execution time for small cases (< 100 qubits) and 50 s for larger cases (> 100 qubits). The eigenstate with the lowest associated eigenvalue was chosen to represent the result of the simulation.

#### Qiskit Qasm Simulator

The BQM was implemented in qiskit through the IBM Experience web interface. This interface provides access to IBM Q hardware and an integrated developer environment served by Jupyter Notebooks with all fundamental Qiskit 0.16.4 python libraries installed.^41^ The IBM Experience provides limited free access to quantum devices, but the available devices were too small (<= 15 qubits) to sample the codon optimization BQM problem, so a noisy simulator hosted by the IBM Experience on classical hardware was used in its place.

The program was constructed and executed using python libraries provided by IBM Qiskit. The core of the implementation builds on the QuadraticProgram base class from the qiskit.optimization library. The codons were appended to the model as binary variables, physically represented by qubits. The objective function was constructed as a minimization with the one- and two-body precomputed terms and an offset of 0.0. The model was simulated using the Aer class with the qasm_simulator backend and FakeVigo noise data, both from the Qiskit library, and the simulator was converted to an executable Quadratic Unconstrained Binary Optimization (QUBO) model with the MinimumEigenOptimizer from the qiskit.optimization.algorithms library. Finally, the combinatorial optimization was carried out with QAOA^37^ from the qiskit.aqua.algorithms library. The program was run with default values where possible.

#### Classic genetic algorithm

A basic single-threaded genetic algorithm with an objective function mathematically equivalent to equation (15) was implemented in Python 3.7 using standard libraries and BioPython 1.78.^47^ For polypeptide fragments (<= 20 residues), the simulations were run for 100 iterations (generations), with each generation procreating 50 times. For full-length sequences (> 100 residues), the number of iterations was increased to 6000. The calculations were run on the Leap web interface to provide an accurate execution time comparison to the AQC calculations. See the Genetic Algorithm Validation section in the Supplementary Information for performance metrics.

#### Test applications

Sample peptide sequences were obtained by splitting the human severe acute respiratory syndrome coronavirus 2 (2019-nCoV, SARS-CoV-2) spike glycoprotein sequence (UniProtKB–P0DTC2) into shorter peptide fragments for simulation on the resource-constrained systems. The sample preparation for the D-Wave system yielded 62 peptides of length 20 and one additional peptide of length 15. The IBM Qasm simulator was simulated with 313 peptides of length 4 and one additional peptide of length 3. Additionally, 10 full-length proteins associated with SARS-CoV-2 studies were scored using the AQC and the GA for comparison. See the Sequences section of the Supplementary Information for the full list.

## Acknowledgements

We acknowledge the use of IBM Quantum services for this work. The views expressed are those of the authors, and do not reflect the official policy or position of IBM or the IBM Quantum team. We would like to thank Eric Manas, Chris MacDermaid and Simon Kelow of GSK for evaluation and suggested refinements, and Andrea Schreij for helping with graphic design.

## Author Contributions

D.M.F. conceived, designed, and implemented this project. R.C.W. contributed to the design of experiments. R.C.W. and K.M.M. contributed to the validation of the work.

D.M.F. and R.C.W. wrote the manuscript.

## Footnotes

1 https://www.dwavesys.com/sites/default/files/Advantage_Datasheet_v9_0.pdf

