## Supplemental Information for "mRNA codon optimization on quantum computers"

### Supplementary Information

#### GC-content derivation

We use the following objective function to measure optimality of GC-content:

$$\Delta = (\rho_{GC} - \rho_T)^2 \quad (S1),$$

where $\rho_T \epsilon\ [0,1] \subset \mathbb{R}$ is the target GC-content,

$$\rho_{GC} = \frac{1}{N}\sum_{i}^{N} s_i q_i \quad (S2)$$

is the GC-content of the nucleotide sequence, $q_i \in \{0,1\}$ denotes the value of qubit *i*, and $s_i$ is the calculation of the normalized GC-content for each codon. Equation (S1) is rewritten as a two-body Hamiltonian, $\mathcal{H}_{GC}$, by expanding the terms and substituting the definition of $\rho_{GC}$ from equation (S2):

$$\begin{aligned}\mathcal{H}_{GC} \propto (\rho_{GC} - \rho_T)^2 &= \rho_{GC}^2 - 2\rho_{GC}\rho_T + \rho_T^2 \\ &= \left(\frac{1}{N}\sum_{i}^{N} s_i q_i\right)^2 - 2\rho_T \frac{1}{N}\sum_{i}^{N} s_i q_i + \rho_T^2 \\ &= \frac{1}{N^2}\left(\sum_{i}^{N} s_i q_i\right)\left(\sum_{j}^{N} s_j q_j\right) - \frac{2\rho_T}{N}\sum_{i}^{N} s_i q_i + \rho_T^2 \quad (S3).\end{aligned}$$

The two terms in parentheses in equation (S3) can be rewritten as an outer product. To see this, explicitly write out the terms in the sums and multiply the terms in parentheses:

$$= \frac{1}{N^2}(s_0q_0 + s_1q_1 + \cdots)(s_0q_0 + s_1q_1 + \cdots) - \frac{2\rho_T}{N}\sum_i^N s_iq_i + \rho_T^2$$

$$= \frac{1}{N^2}(s_0q_0s_0q_0 + s_1q_1s_0q_0 + s_1q_1s_0q_0 \ldots) - \frac{2\rho_T}{N}\sum_i^N s_iq_i + \rho_T^2$$

The terms in parentheses include all combinations of indices, which is compactly written as an outer product:

$$= \frac{1}{N^2}\sum_i^N\sum_j^N (s\otimes s)_{ij}q_iq_j - \frac{2\rho_T}{N}\sum_i^N s_iq_i + \rho_T^2 \quad (S4)$$

Note, this expression strongly resembles the gyration tensor which could be leveraged to compute properties describing the distribution of G's and C's in the sequence. The double sum needs to be restricted to only include terms from the upper triangular elements of the matrix. This is achieved by recognizing the symmetry in matrix; the lower triangular and upper triangular elements are equal, so the sum over the upper triangular terms multiplied by two accounts for all off-diagonal terms.

$$\sum_i^N\sum_{j\neq i}^N (s\otimes s)_{ij}q_iq_j) \; = 2\sum_i^N\sum_{j<\mathrm{i}}^N s_is_jq_iq_j + \sum_i^N s_i^2q_i$$

The sum over the diagonal elements is called the trace, and can be written as a single sum:

$$Tr\left[\sum_i^N\sum_j^N (s\otimes s)_{ij}q_iq_j)\right] = \sum_i^N s_i^2q_i$$

Note, because $q_i$ is restricted to 0 or 1, it is idempotent with itself, so $q_i*q_i = q_i$. A tunable constant is introduced to set the relative importance of this term to compared others. Thus equation (15) from the main text is recovered:

$$\mathcal{H}_{GC} = 2c_{GC}\sum_i^N\sum_{j<i}^N (s\otimes s)_{ij}q_iq_j + c_{GC}\sum_i^N s_i^2q_i - 2\rho_Tc_{GC}\sum_i^N s_iq_i + c_{GC}\rho_T^2 \quad (S5)$$

### BQM challenges and limitations

Mapping codon optimization to a BQM introduces invalid states to the solution space. A set of constraints are introduced to limit the probability of accessing these invalid states, but quantum devices are subject to noise and therefore invalid states cannot be avoided altogether. Let $S_c$ represent the size of the valid solution space, and let

$$\vec{n} = (n_0, \ldots, n_N) \quad (S6)$$

represent the number of possible codons for each position in a polypeptide sequence of length *N*. The size of the solution space is given by the product of the elements of $\vec{n}$:

$$S_c = \prod_i^N n_i \quad (S7).$$

The formulation of the BQM maps every codon that can map to the polypeptide sequence to a qubit, which can be read as either a "0" or a "1". The total size of the BQM solution space, $S_q$, is then given by:

$$S_q = 2^{\sum_i^N n_i} \quad (S8).$$

Rewriting equation (S8) as a product yields:

$$S_q = \prod_i^N 2^{n_i} \quad (S9).$$

Thus, the ratio of valid states to invalid states in the BQM, $f_v$, decreases exponentially as a function of *N*:

$$f_v = \frac{S_c}{S_q} = \prod_i^N n_i 2^{-n_i} \quad (S10).$$

To illustrate the consequences of this expanded solution space, consider a sequence containing 10 methionine residues, which each map to one codon. Then the total solution space only contains 1 valid solution:

$$S_c(MMMMMMMMMM) = \prod_i^N 1 = 1.$$

However, the BQM solution space contains many results:

$$S_q(MMMMMMMMMM) = \prod_i^N 2^1 = 2^{10}.$$

In this case, the BQM is faced with finding the one valid solution in a space containing $2^{10}$ possibilities. Now consider a special case in which each residue in a sequence of length $N$ maps to 4 codons, which is approximately equal to the weighted average of amino acid to codon mappings. In this case,

$$f_v(4\ codons\ per\ position) = \frac{S_c}{S_q} = \frac{\prod_i^N 4}{\prod_i^N 2^4} = \frac{2^{2N}}{2^{4N}} = \frac{1}{2^{2N}} = \frac{1}{S_c}.$$

For each valid state in the BQM, there are $2^{2N}$ invalid states, which is equal in size to the valid solution space itself. In practice, this means perturbations due to noise are highly likely to push the system into an invalid state which can be difficult to recover from since there are exponentially more invalid states compared to valid states.

### Genetic Algorithm Validation

The GA was run on 72 peptide fragments derived from A0A2U1LIM9 of length 10 and the lowest value was compared to the true ground state by exhaustively scoring all possible nucleotide sequences in the solution space. The number of iterations required to identify the global minimum was recorded for each sequence, and the data was plotted as a histogram in Supplementary Figure S3. The maximum number of required steps to reach convergence was 19. Small peptide fragments were therefore each run 20 times for 100 iterations to ensure that the global minimum was identified for each case.

The number of required iterations increases as a function of sequence length. To heuristically determine the number of iterations required to asymptotically converge full-length sequences (100-1,000 amino acids), protein A0A2U1LIM9 (the longest sequence in the test set) was simulated 10 times with 15,000 iterations each (Supplementary Figure S4). The calculation was determined to asymptotically converge within 6,000 iterations.

### Supplementary Figures

| | $q_0$ | $q_1$ | $q_2$ | $q_3$ | $q_4$ | $q_5$ | $q_6$ | $q_7$ | $q_8$ | $q_9$ | $q_{10}$ | $q_{11}$ |
|---|---|---|---|---|---|---|---|---|---|---|---|---|
| $q_0$ | 0 | 0 | 0 | 0 | 3 | 3 | 3 | 3 | 3 | 3 | 0 | 0 |
| $q_1$ | 0 | 0 | 0 | 0 | 3 | 3 | 3 | 3 | 3 | 3 | 0 | 0 |
| $q_2$ | 0 | 0 | 0 | 0 | 3 | 3 | 3 | 3 | 3 | 3 | 0 | 0 |
| $q_3$ | 0 | 0 | 0 | 0 | 8 | 8 | 8 | 8 | 8 | 8 | 0 | 0 |
| $q_4$ | 3 | 3 | 3 | 8 | 0 | 0 | 0 | 0 | 0 | 0 | 8 | 3 |
| $q_5$ | 3 | 3 | 3 | 8 | 0 | 0 | 0 | 0 | 0 | 0 | 8 | 3 |
| $q_6$ | 3 | 3 | 3 | 8 | 0 | 0 | 0 | 0 | 0 | 0 | 15 | 8 |
| $q_7$ | 3 | 3 | 3 | 8 | 0 | 0 | 0 | 0 | 0 | 0 | 8 | 3 |
| $q_8$ | 3 | 3 | 3 | 8 | 0 | 0 | 0 | 0 | 0 | 0 | 8 | 3 |
| $q_9$ | 3 | 3 | 3 | 8 | 0 | 0 | 0 | 0 | 0 | 0 | 8 | 3 |
| $q_{10}$ | 0 | 0 | 0 | 0 | 8 | 8 | 15 | 8 | 8 | 8 | 0 | 0 |
| $q_{11}$ | 0 | 0 | 0 | 0 | 3 | 3 | 8 | 3 | 3 | 3 | 0 | 0 |

Supplementary Figure S1. Result of applying equation (9) from the main text to the example system shown in Figure 1 from the main text. Positions highlighted in yellow represent couplings between the first and second codon, and lavender represents couplings between the second and third codons.

| | $q_0$ | $q_1$ | $q_2$ | $q_3$ | $q_4$ | $q_5$ | $q_6$ | $q_7$ | $q_8$ | $q_9$ | $q_{10}$ | $q_{11}$ |
|---|---|---|---|---|---|---|---|---|---|---|---|---|
| $q_0$ | 0 | $\infty$ | $\infty$ | $\infty$ | 0 | 0 | 0 | 0 | 0 | 0 | 0 | 0 |
| $q_1$ | $\infty$ | 0 | $\infty$ | $\infty$ | 0 | 0 | 0 | 0 | 0 | 0 | 0 | 0 |
| $q_2$ | $\infty$ | $\infty$ | 0 | $\infty$ | 0 | 0 | 0 | 0 | 0 | 0 | 0 | 0 |
| $q_3$ | $\infty$ | $\infty$ | $\infty$ | 0 | 0 | 0 | 0 | 0 | 0 | 0 | 0 | 0 |
| $q_4$ | 0 | 0 | 0 | 0 | 0 | $\infty$ | $\infty$ | $\infty$ | $\infty$ | $\infty$ | 0 | 0 |
| $q_5$ | 0 | 0 | 0 | 0 | $\infty$ | 0 | $\infty$ | $\infty$ | $\infty$ | $\infty$ | 0 | 0 |
| $q_6$ | 0 | 0 | 0 | 0 | $\infty$ | $\infty$ | 0 | $\infty$ | $\infty$ | $\infty$ | 0 | 0 |
| $q_7$ | 0 | 0 | 0 | 0 | $\infty$ | $\infty$ | $\infty$ | 0 | $\infty$ | $\infty$ | 0 | 0 |
| $q_8$ | 0 | 0 | 0 | 0 | $\infty$ | $\infty$ | $\infty$ | $\infty$ | 0 | $\infty$ | 0 | 0 |
| $q_9$ | 0 | 0 | 0 | 0 | $\infty$ | $\infty$ | $\infty$ | $\infty$ | $\infty$ | 0 | 0 | 0 |
| $q_{10}$ | 0 | 0 | 0 | 0 | 0 | 0 | 0 | 0 | 0 | 0 | 0 | $\infty$ |
| $q_{11}$ | 0 | 0 | 0 | 0 | 0 | 0 | 0 | 0 | 0 | 0 | $\infty$ | 0 |

Supplementary Figure S2. Application of $\delta'$ function (equation (12) from main text) to the example system shown in Figure 1 from the main text. The codons mapping to the same sequence position are highlighted in orange, green, and blue for positions 1, 2, and 3, respectively.

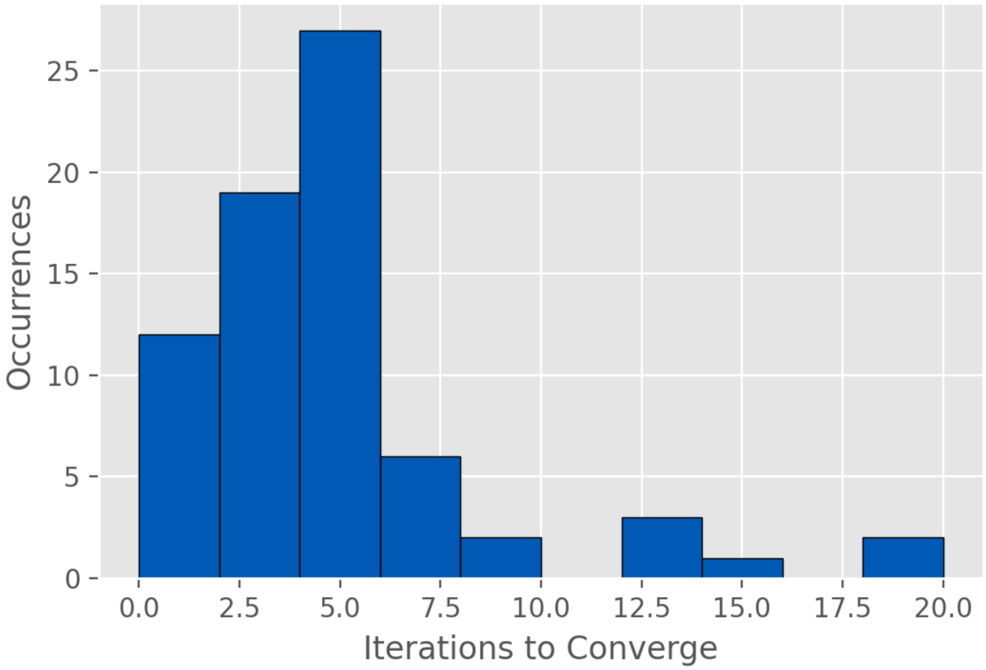


Supplementary Figure S3. Required number of iterations for 72 peptide fragments of length 10 from A0A2U1LIM9 to converge. Convergence was verified by exhaustively enumerating all possible nucleotide sequences and saving the lowest score.

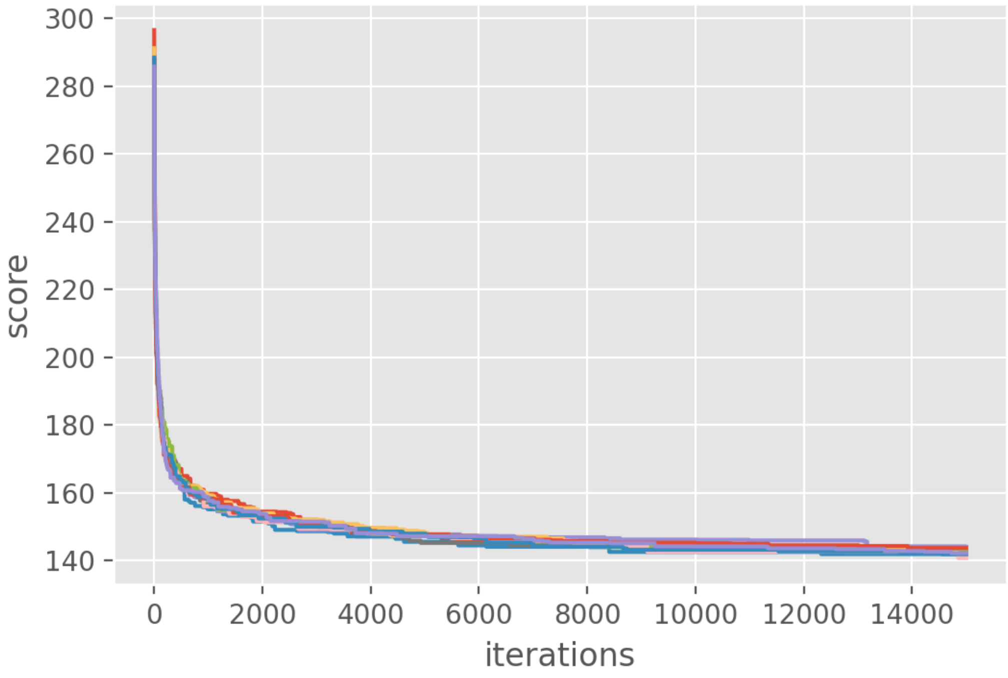


Supplementary Figure S4. Score vs number of GA iterations for protein sequence A0A2U1LIM9, which has 704 amino acids.

### Protein Sequences

```
>sp|P0DTC2|SPIKE_SARS2 Spike glycoprotein OS=Severe acute respiratory syndrome coronavirus 2 OX=2697049
GN=S PE=1 SV=1
MFVFLVLLPLVSSQCVNLTTRTQLPPAYTNSFTRGVYYPDKVFRSSVLHSTQDLFLPFFS
NVTWFHAIHVSGTNGTKRFDNPVLPFNDGVYFASTEKSNIIRGWIFGTTLDSKTQSLLIV
NNATNVVIKVCEFQFCNDPFLGVYYHKNNKSWMESEFRVYSSANNCTFEYVSQPFLMDLE
GKQGNFKNLREFVFKNIDGYFKIYSKHTPINLVRDLPQGFSALEPLVDLPIGINITRFQT
LLALHRSYLTPGDSSSGWTAGAAAYYVGYLQPRTFLLKYNENGTITDAVDCALDPLSETK
CTLKSFTVEKGIYQTSNFRVQPTESIVRFPNITNLCPFGEVFNATRFASVYAWNRKRISN
CVADYSVLYNSASFSTFKCYGVSPTKLNDLCFTNVYADSFVIRGDEVRQIAPGQTGKIAD
YNYKLPDDFTGCVIAWNSNNLDSKVGGNYNYLYRLFRKSNLKPFERDISTEIYQAGSTPC
NGVEGFNCYFPLQSYGFQPTNGVGYQPYRVVVLSFELLHAPATVCGPKKSTNLVKNKCVN
FNFNGLTGTGVLTESNKKFLPFQQFGRDIADTTDAVRDPQTLEILDITPCSFGGVSVITP
GTNTSNQVAVLYQDVNCTEVPVAIHADQLTPTWRVYSTGSNVFQTRAGCLIGAEHVNNSY
ECDIPIGAGICASYQTQTNSPRRARSVASQSIIAYTMSLGAENSVAYSNNSIAIPTNFTI
SVTTEILPVSMTKTSVDCTMYICGDSTECSNLLLQYGSFCTQLNRALTGIAVEQDKNTQE
VFAQVKQIYKTPPIKDFGGFNFSQILPDPSKPSKRSFIEDLLFNKVTLADAGFIKQYGDC
LGDIAARDLICAQKFNGLTVLPPLLTDEMIAQYTSALLAGTITSGWTFGAGAALQIPFAM
QMAYRFNGIGVTQNVLYENQKLIANQFNSAIGKIQDSLSSTASALGKLQDVVNQNAQALN
TLVKQLSSNFGAISSVLNDILSRLDKVEAEVQIDRLITGRLQSLQTYVTQQLIRAAEIRA
SANLAATKMSECVLGQSKRVDFCGKGYHLMSFPQSAPHGVVFLHVTYVPAQEKNFTTAPA
ICHDGKAHFPREGVFVSNGTHWFVTQRNFYEPQIITTDNTFVSGNCDVVIGIVNNTVYDP
LQPELDSFKEELDKYFKNHTSPDVDLGDISGINASVVNIQKEIDRLNEVAKNLNESLIDL
QELGKYEQYIKWPWYIWLGFIAGLIAIVMVTIMLCCMTSCCSCLKGCCSCGSCCKFDEDD
SEPVLKGVKLHYT
>sp|P07711|CATL1_HUMAN Procathepsin L OS=Homo sapiens OX=9606 GN=CTSL PE=1 SV=2
MNPTLILAAFCLGIASATLTFDHSLEAQWTKWKAMHNRLYGMNEEGWRRAVWEKNMKMIE
LHNQEYREGKHSFTMAMNAFGDMTSEEFRQVMNGFQNRKPRKGKVFQEPLFYEAPRSVDW
REKGYVTPVKNQGQCGSCWAFSATGALEGQMFRKTGRLISLSEQNLVDCSGPQGNEGCNG
GLMDYAFQYVQDNGGLDSEESYPYEATEESCKYNPKYSVANDTGFVDIPKQEKALMKAVA
TVGPISVAIDAGHESFLFYKEGIYFEPDCSSEDMDHGVLVVGYGFESTESDNNKYWLVKN
SWGEEWGMGGYVKMAKDRRNHCGIASAASYPTV
>sp|P49350|FPPS_ARTAN Farnesyl pyrophosphate synthase OS=Artemisia annua OX=35608 GN=FPS1 PE=1 SV=2
MSSIDLKSKFLKVYDTLKSELINDPAFEFDDDSRQWIEKMLDYNVPGGKLNRGLSVVDSY
QLLKGGELSDDEIFLSSALGWCIEWLQAYFLVLDDIMDESHTRRGQPCWFRLPKVGMIAA
NDGILLRNHVPRILKKHFRGKPYYVDLVDLFNEVEFQTASGQMIDLITTLVGEKDLSKYS
LSIHRRIVQYKTAYYSFYLPVACALLMFGEDLDKHVEVKNVLVEMGTYFQVQDDYLDCFG
APEVIGKIGTDIEDFKCSWLVVKALELANEEQKKVLHENYGKKDPASVAKVKEVYHTLNL
QAVFEDYEATSYKKLITSIENHPSKAVQAVLKSFLGKIYKRQK
>sp|A0A2U1Q018|ADH1_ARTAN Alcohol dehydrogenase 1 OS=Artemisia annua OX=35608 GN=ADH1 PE=1 SV=1
MAQKAPGVITCKAAVVWELGGPVVLEEIRVDPPKASEVRIKMLCASLCHTDVLCTKGFPI
PLFPRIPGHEGVGVIESVGKDAKGLKPGDIVMPLYLGECGQCLNCKTGKTNLCHVYPPSF
SGLMNDGTSRMSIARTGESIYHFASCSTWTEYAVADCNYVLKINPKISYPHASFLSCGFT
TGFGATWRETQVSKGSSVAVFGIGTVGLGVIKGAQLQGASKIIGVDVNQYKAAKGKVFGM
TDFINPKDHPDKSVSELVKELTHGLGVDHCFECTGVPSLLNEALEASKIGIGTVVPIGAG
GEASVAINSLILFSGRTLKFTAFGGVRTQSDLPVIIDKCLNKEIQLDELLTHEIHLDNIQ
EAFEILKKPDCVKILIKF
>sp|P35613-2|BASI_HUMAN Isoform 2 of Basigin OS=Homo sapiens OX=9606 GN=BSG
MAAALFVLLGFALLGTHGASGAAGTVFTTVEDLGSKILLTCSLNDSATEVTGHRWLKGGV
VLKEDALPGQKTEFKVDSDDQWGEYSCVFLPEPMGTANIQLHGPPRVKAVKSSEHINEGE
TAMLVCKSESVPPVTDWAWYKITDSEDKALMNGSESRFFVSSSQGRSELHIENLNMEADP
GQYRCNGTSSKGSDQAIITLRVRSHLAALWPFLGIVAEVLVLVTIIFIYEKRRKPEDVLD
DDDAGSAPLKSSGQHQNDKGKNVRQRNSS
>sp|Q1PS23|AMO_ARTAN Amorpha-4,11-diene 12-monooxygenase OS=Artemisia annua OX=35608 GN=CYP71AV1
PE=1 SV=2
MKSILKAMALSLTTSIALATILLFVYKFATRSKSTKKSLPEPWRLPIIGHMHHLIGTTPH
RGVRDLARKYGSLMHLQLGEVPTIVVSSPKWAKEILTTYDISFANRPETLTGEIVLYHNT
DVVLAPYGEYWRQLRKICTLELLSVKKVKSFQSLREEECWNLVQEIKASGSGRPVNLSEN
VFKLIATILSRAAFGKGIKDQKELTEIVKEILRQTGGFDVADIFPSKKFLHHLSGKRARL
TSLRKKIDNLIDNLVAEHTVNTSSKTNETLLDVLLRLKDSAEFPLTSDNIKAIILDMFGA
GTDTSSSTIEWAISELIKCPKAMEKVQAELRKALNGKEKIHEEDIQELSYLNMVIKETLR
LHPPLPLVLPRECRQPVNLAGYNIPNKTKLIVNVFAINRDPEYWKDAEAFIPERFENSSA
TVMGAEYEYLPFGAGRRMCPGAALGLANVQLPLANILYHFNWKLPNGVSYDQIDMTESSG
ATMQRKTELLLVPSF
>sp|A0A2U1LIM9|NCPR1_ARTAN NADPH--cytochrome P450 reductase 1 OS=Artemisia annua OX=35608 GN=CPR1
PE=1 SV=2
MQSTTSVKLSPFDLMTALLNGKVSFDTSNTSDTNIPLAVFMENRELLMILTTSVAVLIGC
VVVLVWRRSSSAAKKAAESPVIVVPKKVTEDEVDDGRKKVTVFFGTQTGTAEGFAKALVE
EAKARYEKAVFKVIDLDDYAAEDDEYEEKLKKESLAFFFLATYGDGEPTDNAARFYKWFT
EGEEKGEWLEKLQYAVFGLGNRQYEHFNKIAKVVDEKLVEQGAKRLVPVGMGDDDQCIED
DFTAWKELVWPELDQLLRDEDDTSVATPYTAAVAEYRVVFHDKPETYDQDQLTNGHAVHD
AQHPCRSNVAVKKELHSPLSDRSCTHLEFDISNTGLSYETGDHVGVYVENLSEVVDEAEK
LIGLPPHTYFSVHTDNEDGTPLGGASLPPPFPPCTLRKALASYADVLSSPKKSALLALAA
HATDSTEADRLKFLASPAGKDEYAQWIVASHRSLLEVMEAFPSAKPPLGVFFASVAPRLQ
```

```
PRYYSISSSPKFAPNRIHVTCALVYEQTPSGRVHKGVCSTWMKNAVPMTESQDCSWAPIY
VRTSNFRLPSDPKVPVIMIGPGTGLAPFRGFLQERLAQKEAGTELGTAILFFGCRNRKVD
FIYEDELNNFVETGALSELVTAFSREGATKEYVQHKMTQKASDIWNLLSEGAYLYVCGDA
KGMAKDVHRTLHTIVQEQGSLDSSKAELYVKNLQMAGRYLRDVW
>sp|P0DTC9|NCAP_SARS2 Nucleoprotein OS=Severe acute respiratory syndrome coronavirus 2 OX=2697049
GN=N PE=1 SV=1
MSDNGPQNQRNAPRITFGGPSDSTGSNQNGERSGARSKQRRPQGLPNNTASWFTALTQHG
KEDLKFPRGQGVPINTNSSPDDQIGYYRRATRRIRGGDGKMKDLSPRWYFYYLGTGPEAG
LPYGANKDGIIWVATEGALNTPKDHIGTRNPANNAAIVLQLPQGTTLPKGFYAEGSRGGS
QASSRSSSRSRNSSRNSTPGSSRGTSPARMAGNGGDAALALLLLDRLNQLESKMSGKGQQ
QQGQTVTKKSAAEASKKPRQKRTATKAYNVTQAFGRRGPEQTQGNFGDQELIRQGTDYKH
WPQIAQFAPSASAFFGMSRIGMEVTPSGTWLTYTGAIKLDDKDPNFKDQVILLNKHIDAY
KTFPPTEPKKDKKKKADETQALPQRQKKQQTVTLLPAADLDDFSKQLQQSMSSADSTQA
>sp|P0DTC3|AP3A_SARS2 ORF3a protein OS=Severe acute respiratory syndrome coronavirus 2 OX=2697049
GN=3a PE=1 SV=1
MDLFMRIFTIGTVTLKQGEIKDATPSDFVRATATIPIQASLPFGWLIVGVALLAVFQSAS
KIITLKKRWQLALSKGVHFVCNLLLLFVTVYSHLLLVAAGLEAPFLYLYALVYFLQSINF
VRIIMRLWLCWKCRSKNPLLYDANYFLCWHTNCYDYCIPYNSVTSSIVITSGDGTTSPIS
EHDYQIGGYTEKWESGVKDCVVLHSYFTSDYYQLYSTQLSTDTGVEHVTFFIYNKIVDEP
EEHVQIHTIDGSSGVVNPVMEPIYDEPTTTTSVPL
>sp|P0DTC5|VME1_SARS2 Membrane protein OS=Severe acute respiratory syndrome coronavirus 2
OX=2697049 GN=M PE=3 SV=1
MADSNGTITVEELKKLLEQWNLVIGFLFLTWICLLQFAYANRNRFLYIIKLIFLWLLWPV
TLACFVLAAVYRINWITGGIAIAMACLVGLMWLSYFIASFRLFARTRSMWSFNPETNILL
NVPLHGTILTRPLLESELVIGAVILRGHLRIAGHHLGRCDIKDLPKEITVATSRTLSYYK
LGASQRVAGDSGFAAYSRYRIGNYKLNTDHSSSSDNIALLVQ
>sp|P0DTC7|NS7A_SARS2 ORF7a protein OS=Severe acute respiratory syndrome coronavirus 2 OX=2697049
GN=7a PE=1 SV=1
MKIILFLALITLATCELYHYQECVRGTTVLLKEPCSSGTYEGNSPFHPLADNKFALTCFS
TQFAFACPDGVKHVYQLRARSVSPKLFIRQEEVQELYSPIFLIVAAIVFITLCFTLKRKT
E
```
